# Maternal exercise alters placental proteome in an exercise mode-specific manner

**DOI:** 10.1101/2024.11.11.623023

**Authors:** Filip Jevtovic, Breanna Wisseman, Fahmida Jahan, Alex Claiborne, David N. Collier, James E. DeVente, Steven Mouro, Tonya Zeczycki, Laurie J. Goodyear, Linda E. May

## Abstract

Maternal exercise is a widely recommended and safe intervention associated with the improvement of maternal gestational and infant metabolic health. While various modes of exercise are deemed safe during pregnancy, the effects of supervised maternal aerobic, resistance, and combination (aerobic+resistance) exercise remain understudied. Specifically, it remains unknown how different modes of maternal exercise affect the placenta, an organ central to maternal-fetal communication and successful pregnancy outcomes. This study aimed to characterize the placental proteomic changes in response to controlled and supervised maternal exercise during gestation. Results showed that the placental proteomic landscape changes in an maternal exercise mode-specific way. Additionally, proteomics revealed that ∼20% of the identified placental proteins were associated with maternal exercise volume during gestation. These results highlight the differential effect maternal exercise modes have on the placental proteome and further implicate the placenta in mediating the effects of maternal exercise on maternal and infant health.

## Introduction

Maternal exercise has been established as a safe non-pharmacological intervention to improve maternal gestational health and pregnancy outcomes[1–7]. Furthermore, maternal exercise has been associated with improvements in infant anthropometrics [8–10], whole-body[11] and cellular metabolism[8,12–14], infant cardiovascular health[15–17], and neuromotor development[18]. Nevertheless, despite the profound benefits of ME, the mechanisms that mediate maternal gestational, pregnancy, and infant health remain to be understood.

Central to maternal-fetal communication, the placenta is a heterogeneous organ that displays prominent plasticity to environmental stimuli experienced by the mother. The placenta plays a pivotal role in maternal health maintenance and the transmission of environmental inputs to the growing fetus. Thus far, placental clinical research is predominantly disease-centric with significant gaps in describing the effects of maternal exercise on the human placenta[19]. Further, current maternal exercise research remains observational[20–24], rather than controlled and supervised, and rarely utilizes comprehensive -omics analyses that capture the complexity of placental response to supervised ME. While these shortcomings have been addressed in rodent models (reviewed in [14,25,26]); differences in placental structure and physiology, and the chronology of fetal development prevent direct comparison of the maternal exercise effects between species[27]. Finally, gaps in both human and animal literature exist regarding the effects of supervised maternal exercise modes on the placenta, as research so far is confined to aerobic exercise.

To address these gaps and understand how placenta physiology adapts to supervised maternal exercise modes, we have undertaken a comparative, untargeted proteome analysis of term placentas from women who performed supervised, controlled aerobic, resistance, or combination (aerobic+resistance) moderate-intensity exercise during pregnancy or attention-controls. Since proteins play essential roles in maintaining cellular structure, function, and homeostasis, exploring the unique proteomic landscape of placental tissue provides invaluable insight into placental ‘health’ and biological function, as well as reveal how the placenta adapts to various maternal stimuli. Here, we describe the placental proteomic changes in response to different supervised maternal exercise modes to uncover the molecular underpinnings driving the profound effects of ME. Additionally, it will lay the ground work for additional hypothesis-driven research focusing on the mechanisms of maternal-fetal communication.

## Methods

### Ethics statement

This study used placental tissue collected in the ENHANCED (Enhanced Neonatal Health and Neonatal Cardiac Effect Developmentally) and EMCOR (Pregnancy Exercise Mode Effect on Childhood Obesity) studies (ClinicalTrials.gov Identifiers: NCT03838146 and NCT04805502). Approval for this study and all experiments was obtained from the East Carolina University Institutional Review Board and informed consent was obtained from each participant upon enrollment.

### Pre-intervention testing

Females were recruited between 13-16 weeks of gestation based on the inclusion criteria outlined in the study design (Figure 1). After receiving clearance from their obstetric provider, participants were randomly assigned to aerobic, resistance, combination (aerobic+resistance) exercise, or attention control group.

**Figure 1.**
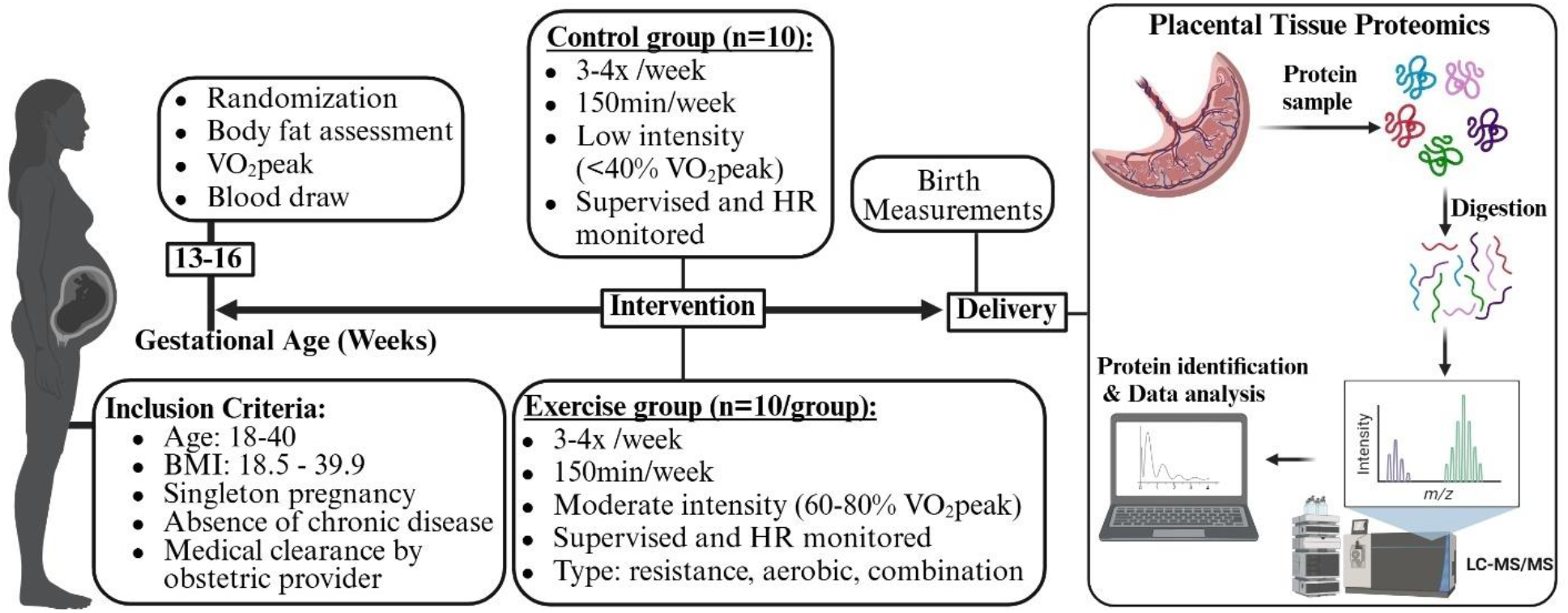
Design and timeline of the study. Created with BioRender.

Participants completed a submaximal modified Balke treadmill test following the previously described submaximal modified Balke treadmill test [28] to measure peak oxygen consumption (VO_2peak_), and to determine target heart rate (THR) zones for subsequent exercise training. THR zones corresponded to maternal HR at 60-80% of maximal oxygen consumption, reflecting moderate intensity [28]. For participants recruited during the COVID-19 pandemic, to minimize exposure and potential risk associated with exercise testing, THR was determined based on the pre-pregnancy physical activity level and age, using published guidelines [28]. Fourteen people had their THR zones estimated based on previously established protocols[28], due to the COVID-19 pandemic; however, placental protein expression was similar between subjects that were assigned exercise intensity based on estimated and measured HR.

### Exercise intervention

As published previously [29], participants exercised according to American College of Obstetricians and Gynecologists guidelines for the duration of their pregnancy (∼16-40 weeks). Participants performed moderate intensity (60%-80% maximal oxygen consumption and 12-14 rated perceived exertion) aerobic, resistance, or a combination exercise. Every session was supervised. Additionally, HR monitors were used to ensure that participants in all exercise groups maintained their HR within their assigned THR zone during each exercise session. Resistance training was done using free weights and seated Cybex machines. Aerobic training was done using treadmills, ellipticals, recumbent bicycles, or stair-stepping equipment. The combination exercise group spent the first half of the session doing resistance training followed by aerobic training. The control group performed supervised stretching, breathing, and flexibility exercises at low intensity (<40% VO2peak). Exercise sessions consisted of a 5-minute warm-up, 50-minutes completing group specific activities, and a 5-minute cool down period. Maternal exercise adherence was calculated by dividing the number of sessions attended by the total number of possible sessions within the participants’ gestational period. Maternal exercise intensity (METs) was based on the published compendium of physical activity for the exercise performed in each session [30]. The average maternal exercise dose during each week, expressed as MET·min/wk, was quantified (frequency X duration of session) and then multiplied by the intensity (METs) of their exercise. Further, the total volume of exercise during pregnancy (Total MET·min) was calculated by multiplying the MET·min/wk by the total number of weeks of gestational exercise. Importantly, average, and total exercise volumes were calculated between 16-36 weeks of gestation, to avoid the influence of different gestational lengths between mothers (37-41 weeks) on exercise volume.

The ability of our protocol to elicit maternal physiological adaptations to exercise has been previously reported [31].

### Maternal and infant birth measurements

Maternal measurements were obtained as previously described[29]. Maternal age, parity, pre-pregnancy weight and height and body mass index (BMI, kg/m^2^), gestational diabetes mellitus status (yes or no), length of gestation, and mode of delivery were abstracted from various sources including pre-screening eligibility and postpartum questionnaires as well as maternal and neonatal electronic health records. At 16 weeks of gestation, we obtained maternal BMI and determined maternal percent body fat via validated skinfold technique and age-adjusted equations[32,33]. Additionally, maternal fingerstick blood was analyzed using Cholestech LDX Analyzer (Alere Inc., Waltham, MA, USA) and point of care Lactate Plus Analyzer (Nova Biomedical, Waltham, MA, USA) to quantify maternal lipids (total cholesterol (TC), triglycerides (TG), HDL, non-HDL, LDL), glucose and lactate. Birth measurements (weight, length, Ponderal Index, abdominal, head and chest circumference, 1 and 5-minute Apgar scores) and infant sex were extracted from neonatal electronic health records.

### Placental tissue collection and proteomics

Term placentas were collected after delivery, and tissue was sampled from central (2-3 cm from umbilical cord insertion) and peripheral (2-3 cm from the edge of the placental disc) areas while avoiding placental vasculature and areas of calcification and necrosis. Tissues were flash-frozen in liquid nitrogen, powdered on dry ice using mortar and pestle, and homogenized in lysis buffer (Cat. #: 43-040; Merck, Darmstadt, Germany) containing protease inhibitor cocktail (complete Tablets Mini, EDTA-free protease inhibitor cocktail tablets, Roche, Switzerland). Samples were homogenized using a stand homogenizer (Ultra-Turrax, IKA) and then centrifuged for 15 minutes at 4 °C, 10,000xg. Sample protein content was determined using a Pierce BCA Protein Assay Kit (Thermo Scientific, Rockford IL, USA).

To prepare mass spectrometry grade peptides, the protein concentration was adjusted to 2 µg/µL with a modified lysis buffer, where the protease inhibitor was removed. Proteins were precipitated from 50 µL of sample (∼100 µg protein) using ice cold methanol (3:1 v/v) at -20 °C and pelleted by centrifugation. The pellets were washed with ice cold methanol twice and allowed to air dry before further use. The PreOmics iST kit (PreOmics GmbH) was used to prepare mass spectrometry grade peptides. Precipitated proteins were resuspended in the provided lysis/denaturing buffer and denatured and alkylated at 95 °C for 10 min. After cooling to room temperature, proteins were digested for 3 hours at 37 °C while shaking (600 rpm). Samples were then loaded onto the provided microcolumns for purification. Peptides were eluted via centrifugation using the provided elution buffer, dried to completeness under a N_2_ stream, and resuspended in loading buffer (98:2 water:acetonitrile; 0.1% formic acid) at a concentration of 0.25 mg/mL.

Peptides were analyzed by nanoLC-MS/MS using an UltiMate 3000 RSLCnano system coupled to a Q Exactive Plus Hybrid Orbitrap mass Spectrometer (ThermoFisher) via nanoelectrospray ionization. Peptides were separated using an effective linear gradient of 4-35% acetonitrile (0.1% formic acid) over 135 min. For data-dependent acquisition, MS spectra were acquired in positive mode. MS1 was performed at a resolution of 70,000 with an AGC target of 2×10^5^ ions and a maximum injection time of 100 ms. MS2 spectra were collected on the top 20 most abundant precursor ions with a charge >1 using an isolation window of 1.5 m/z and fixed first mass of 140 m/z. The normalized collision energy was 30. MS2 spectra were acquired at 17,500 resolutions with a maximum injection time of 60 ms, an AGC target of 1×10^5^. Dynamic exclusion was set to 30 sec.

FragPipe (v 19.1)[34,35] was used for raw data analysis with default search parameters for open and Label Free Quantification-Matching Between Runs (LFQ-MBR) workflows unless otherwise noted. Placenta tissue was searched with biological replicates (n=10) for each exercise group identified. An initial open search against the canonical + isoforms Uniprot *Homo sapiens* reference proteome (UP000005640, accessed 11/2023) was used to identify potential post-translational modifications for inclusion in the LFQ-MBR workflow. Precursor m/z tolerance was set to -150 to 500 Da and fragment tolerance was ± 20 ppm with 3 missed cleavages for Tryp and Lys-C allowed. Peptide spectrum matches (PSMs) were validated using PeptideProphet and results were filtered at the ion, peptide, and protein level with a 1% false discovery rate (FDR). Based on these initial searches, the following variable modifications were included in the LFQ-MBR analysis: oxidation (+15.5995 Da on Met), deamidation (+0.98401 Da on Gln and Asn), and fixed modification carbamidomethyl (+57.025 Da on Cys). For LFQ-MBR analysis, data were searched against the canonical Uniprot *Homo sapiens* reference proteome (UP000005640, accessed 11/2023 and 1/2024). Precursor ion m/z tolerance was ± 20 ppm with 3 missed cleavages for Trypsin/LysC allowed. The search results were filtered by a 1% FDR at the ion, peptide, and protein levels. PSMs were validated using Percolator and label-free quantification was carried out using IonQuant[36] Match between runs FDR rate at the ion level was set to 10% for the top 300 runs. Proteins with >95% probability of ID, >2 unique peptides, and in more than 80% of a sample group (i.e. 7/10 injections) were considered high-confidence IDs and retained for analysis. Intensities were log_2_ transformed and normalized to the median intensities of the sample group. Relative abundances for low sampling proteins were determined via normal distribution in Perseus[37].

### Statistical analysis

Maternal and infant characteristics were compared using one-way ANOVA. For comparison of protein abundances between specific exercise groups and the control group (e.g., aerobic vs control), we used multiple unpaired t-tests with Welch correction for individual variance for each group, with p-value <0.1 for statistical significance. Log_2_ fold-change for potentially biologically relevant differences between groups were determined *a priori* as a Log_2_ fold-change greater than 1.2 or less than -0.8. Pearson’s correlations were used to determine if there were any relationships between exercise variables and placental proteins, with statistical significance set at *p* < 0.05. Statistical analyses were performed using GraphPad Prism version 9.3 (GraphPad Software, San Diego, CA) for Windows.

## Results

### Participant characteristics

Maternal and infant characteristics are presented in Table 1. All exercise groups had higher average weekly (MET•min/wk) and total pregnancy exercise volume (MET•min) compared to the control group. All groups had similar pre-pregnancy BMI, body fat percentage, and blood lipids, glucose, and lactate levels at 16 weeks of gestation. All participants were free of gestational diabetes, based on their 1-hour oral glucose tolerance test values obtained between 22-24 weeks of gestation. There was a similar infant sex distribution across the groups and all infants had similar birth weight, anthropometrics, and APGAR scores.

**Table 1.** Maternal and Infant Characteristics.

| Maternal Characteristics | Control (n=10) | Aerobic (n=10) | Combination (n=10) | Resistance (n=10) | p-value |
| --- | --- | --- | --- | --- | --- |
| Age (yrs) | 31.3±3.6 | 30.7±4.8 | 27.8±4.4 | 32.1±5.6 | n.s. |
| VO<sub>2peak</sub> (ml/kg/min)* | 19.7±2.5 | 24.4±4.6 | 19.7±8.5 | 22.8±5.3 | n.s. |
| Average weekly volume between 16-36 weeks of gestation (MET•min/wk) | 182.7±137.2 | 650±118.7 | 602.7±129.9 | 573.3±100.4 | <0.001 |
| Total exercise volume between 16-36 weeks of gestation (MET•min) | 4571±3682 | 15426±3547 | 13946±2629 | 13088±2466 | <0.001 |
| Pre-pregnancy BMI | 28.9±4.6 | 27.7±5.8 | 27.8±5.6 | 24.5±4.3 | n.s. |
| 16-week BMI<sup>#</sup> | 29.1±4.5 | 28.6±6.4 | 29.3±6.7 | 24.6±4.1 | n.s. |
| 16-week Body Fat (%)<sup>#</sup> | 36.1±3.8 | 36.2±3.1 | 35.7±3.5 | 32.5±3.8 | n.s. |
| 16-week Total Cholesterol (mg/dl)<sup>#</sup> | 180.7±29.1 | 171.7±33.84 | 180.5±33.9 | 178.4±40.2 | n.s. |
| 16-week LDL (mg/dl) | 95.6±25.6 | 87.1±30 | 87.1±31.3 | 96.3±31.5 | n.s. |
| 16-week HDL (mg/dl) <sup>#</sup> | 56.8±18.3 | 61.8±16.6 | 65.3±10.6 | 61.9±11.3 | n.s. |
| 16-week non-HDL (mg/dl)<sup>#</sup> | 121.7±24 | 110.1±32.4 | 115.2±31.4 | 116.3±35.9 | n.s. |
| 16-week Triglycerides (mg/dl)<sup>#</sup> | 110.3±38.5 | 113.4±58.4 | 140.7±75.2 | 100±25.3 | n.s. |
| 16-week Glucose (mg/dl) | 84.3±8.9 | 83.7±7.6 | 79.8±6.2 | 77.8±6.3 | n.s. |
| 16-week Lactate (mmol/l) <sup>#</sup> | 1.1±0.4 | 1.4±0.6 | 1.2±0.7 | 1.2±0.9 | n.s. |
| Gestational length (weeks) | 39.5±1.3 | 40.2±0.8 | 39.9±0.5 | 40±1 | n.s. |
| Parity | 1 (0, 3) | 0 (0, 2) | 0 (0, 1) | 0 (0, 2) | n.s. |
| Mode of delivery (SVD/C-section) | 7/3 | 7/3 | 5/5 | 8/2 | n.s. |
| **Infant Characteristics** | **Control (n=10)** | **Aerobic (n=10)** | **Combination (n=10)** | **Resistance (n=10)** | **p-value** |
| Fetal sex (F/M) | 5/5 | 5/5 | 7/3 | 5/5 | n.s. |
| Birth weight (kg) | 3.74±0.46 | 3.47±0.37 | 3.47±0.3 | 3.47±0.3 | n.s. |
| Birth length (m) <sup>#</sup> | 0.49±0.03 | 0.49±0.02 | 0.50±0.02 | 0.49±0.03 | n.s. |
| Birth BMI<sup>#</sup> | 15.2±1.4 | 14.0±1.1 | 14.0±1.5 | 14.7±2.5 | n.s. |
| Head circumference (m) <sup>#</sup> | 0.35±0.02 | 0.34±0.01 | 0.35±0.01 | 0.35±0.01 | n.s. |
| Chest circumference (m) <sup>#</sup> | 0.34±0.02 | 0.33±0.02 | 0.33±0.01 | 0.34±0.01 | n.s. |
| Abdominal circumference (m) <sup>#</sup> | 0.33±0.02 | 0.30±0.07 | 0.03±0.02 | 0.32±0.01 | n.s. |
| Apgar-1 minute<sup>#</sup> | 8 (2, 9) | 8 (8, 9) | 8 (6, 9) | 8 (7, 9) | n.s. |
| Apgar-5 minute<sup>#</sup> | 9 (8, 9) | 9 (9, 9) | 9 (7, 9) | 9 (9, 9) | n.s. |
| All data expressed as mean ± SD, t-test, p≤0.05. * VO<sub>2peak</sub> n=26/40 due to COVID; <sup>#</sup> Missing 1-3/40 values. SVD = spontaneous vaginal delivery; F= female infant sex; M = male infant sex. One-way ANOVA. |  |  |  |  |  |

### Effects of aerobic exercise on placental proteome

From the placental tissue, we identified 2025 proteins with high-confidence across all groups. For the aerobic group, we observed 471 proteins with statistically significant (p<0.1) abundances when compared to the control group (Figure 2A), with 77 of those proteins showing greater than a 1.2 (Figure 2B, red) or less than -0.8 Log_2_ fold-change (Figure 2B, red) in abundance. We used ShinyGO 8.0 (ShinyGO 0.80 (sdstate.edu)) to identify KEGG pathways that differently abundant proteins correspond to, and we further categorized these 77 differentially abundant proteins based on GO molecular function, biological processes, and cellular components (Figure 2E-G); categories represent the annotation of enrichment of specific proteins, with fold-change representing the magnitude of the pathway enrichment (e.g., number of genes altered / total number of genes in the pathway). Finally, we used a human protein atlas to identify secreted proteins from the 77 differentially abundant proteins between aerobic exercise and control groups (Figure 2D).

**Figure 2.**
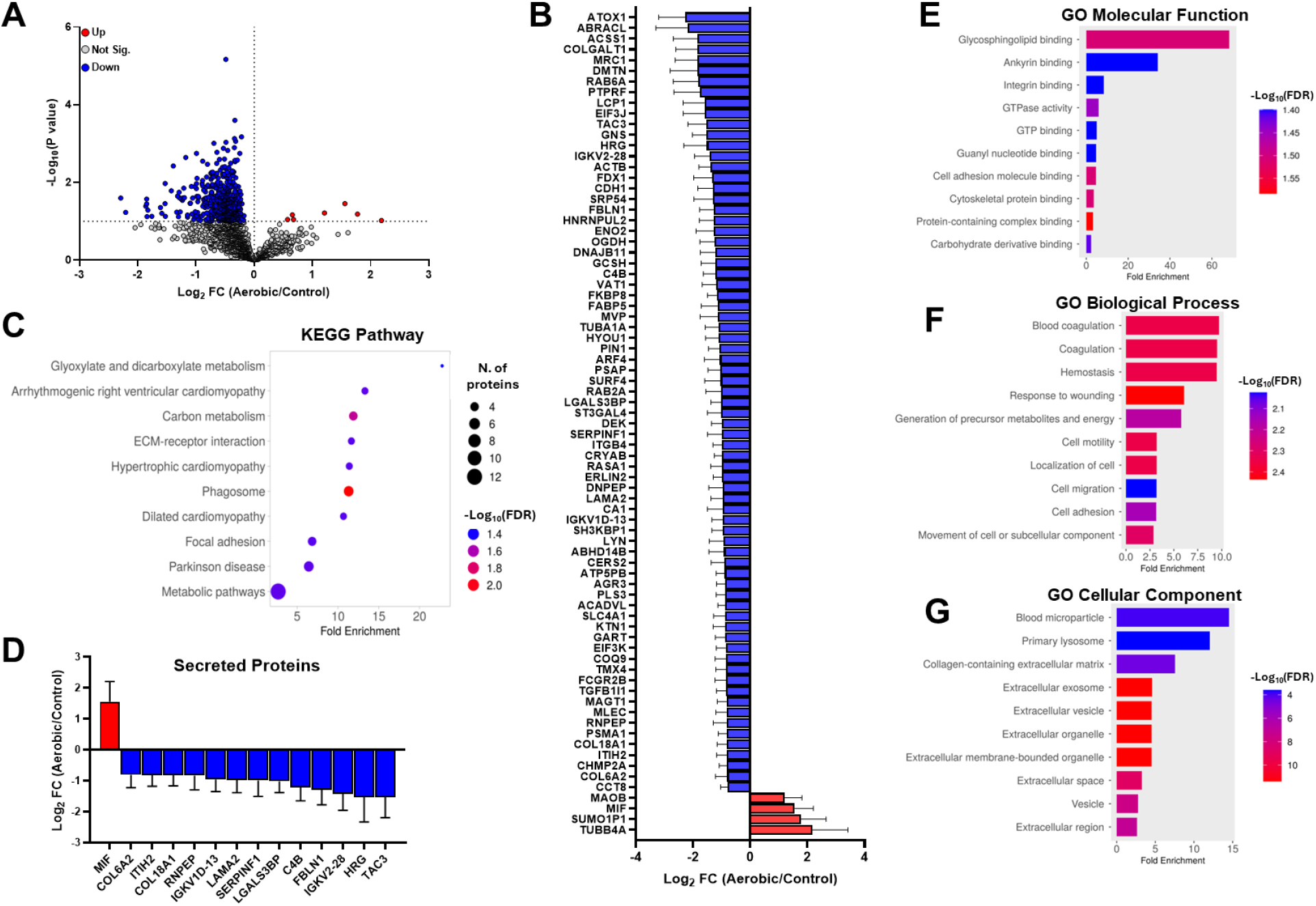
Effects of aerobic exercise on the placental proteome. Volcano plot of Log2 fold-change of protein abundances in aerobic compared to the control groups vs. -Log10 (p-value) for the fold-change. Red symbols indicate proteins with significantly higher abundances in the aerobic vs. control group, blue symbols indicate proteins with significantly lower abundances in the aerobic vs. control group, grey symbols showed no significance in the log2 fold-changes (p>0.1) (A). Plots of the Log2 fold-change for potentially relevant proteins in aerobic compared to control group. Data is shown as the mean of the Log2 fold-change ± SEM (n=10 for each group). Data coloring is the same as in A. (B). KEGG Pathway (C), GO Molecular Function (E), GO Biological Process (F), and GO Cellular Component (G) analysis based on differential protein abundances. Expression of secreted proteins based on the Protein Atlas (D).

### Effects of resistance exercise on placental proteome

In resistance exercise group, we observed 212 proteins with significantly (p<0.01) different abundance when compared to the control group (Figure 3A). 54 of these proteins were within ‘meaningful’ fold change differences (Figure 3B). Further categorizing these proteins based on the KEGG pathway, showcased the involvement of these proteins in complement and coagulation cascades, tight junction, focal adhesion, and regulation of the actin cytoskeleton (Figure 3C). Within 54 differentially abundant proteins we identified 9 secretory proteins that were downregulated with resistance exercise (Figure 3D). Finally, differentially abundant proteins were categorized based on the GO analysis for molecular function, biological processes, and cellular components (Figure 3E-G).

**Figure 3.**
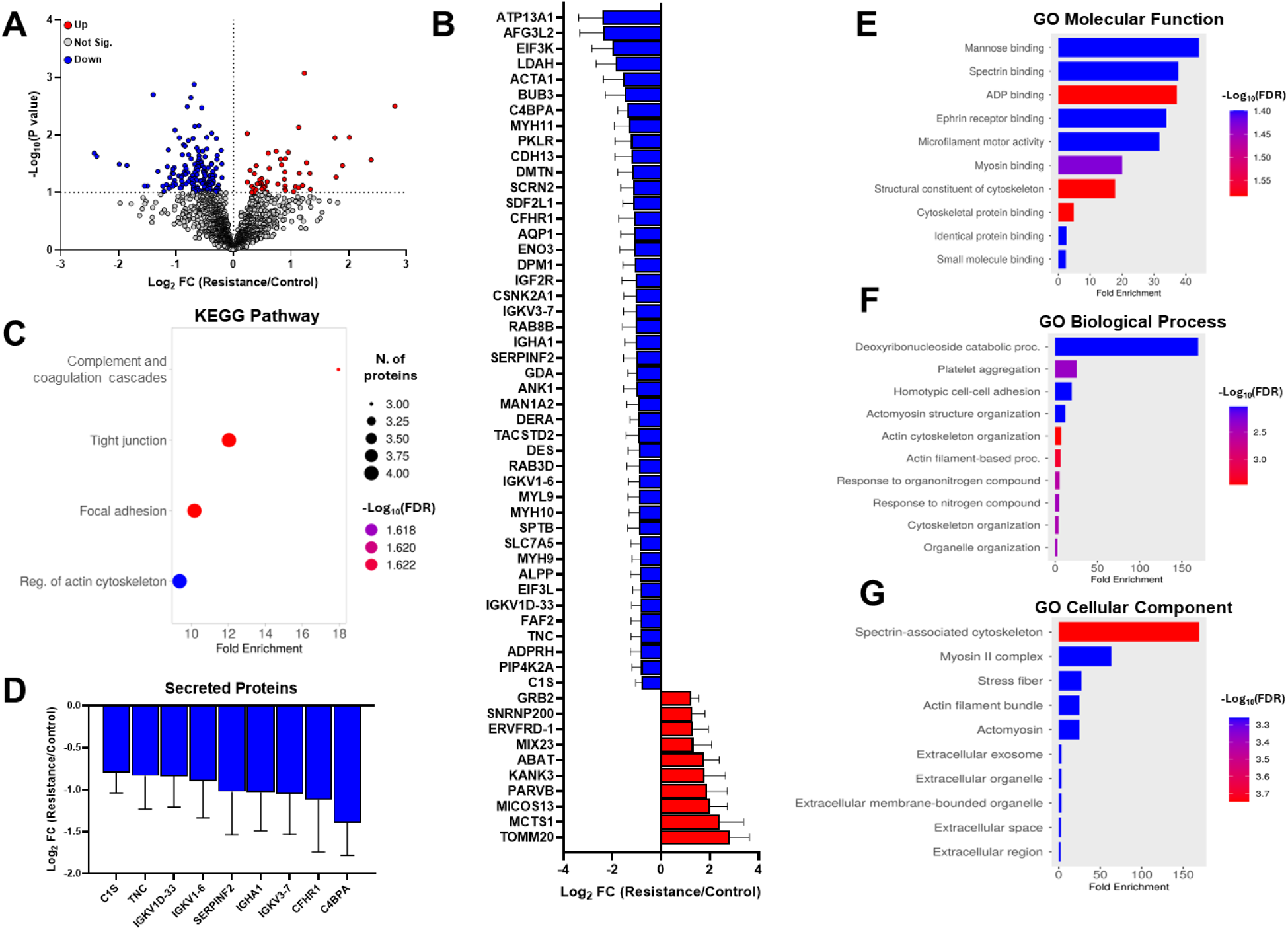
Effects of resistance exercise on the placental proteome. Volcano plot of Log2 fold-changes of protein abundances in resistance vs control group, with -Log10 (p-value) for the fold-change on the y-axis (A). Differentially abundant and significantly up- and downregulated proteins within potentially ‘meaningful’ differences (<-0.8, >1.2) (B). KEGG Pathway (C), GO Molecular Function (E), GO Biological Process (F), and GO Cellular Component (G) based on the differentially abundant proteins with fold-change over 1.2 and less than -0.8. Expression of secreted proteins based on the Protein Atlas (D).

### Effects of combination exercise on placental proteome

Compared to the control group, we identified 104 proteins with significantly (p<0.1) different abundance (Figure 4A). Out of 104, 30 proteins were within the fold-change parameters of >1.2 and <-0.8 (Figure 4B). When put through KEGG pathway analysis, these proteins correspond to butanoate metabolism, and valine leucine and isoleucine pathways (Figure 4C). Furthermore, these 30 proteins correspond to multiple pathways identified in GO molecular function enrichment analysis (Figure 4E); however, identified proteins did not correspond to any pathways in GO Cellular Component. Finally, out of 30 proteins 6 proteins corresponded to the endocytosis pathway based on GO Biological Processes analysis (fold change 8.7; -log10(FDR) 1.37).

**Figure 4.**
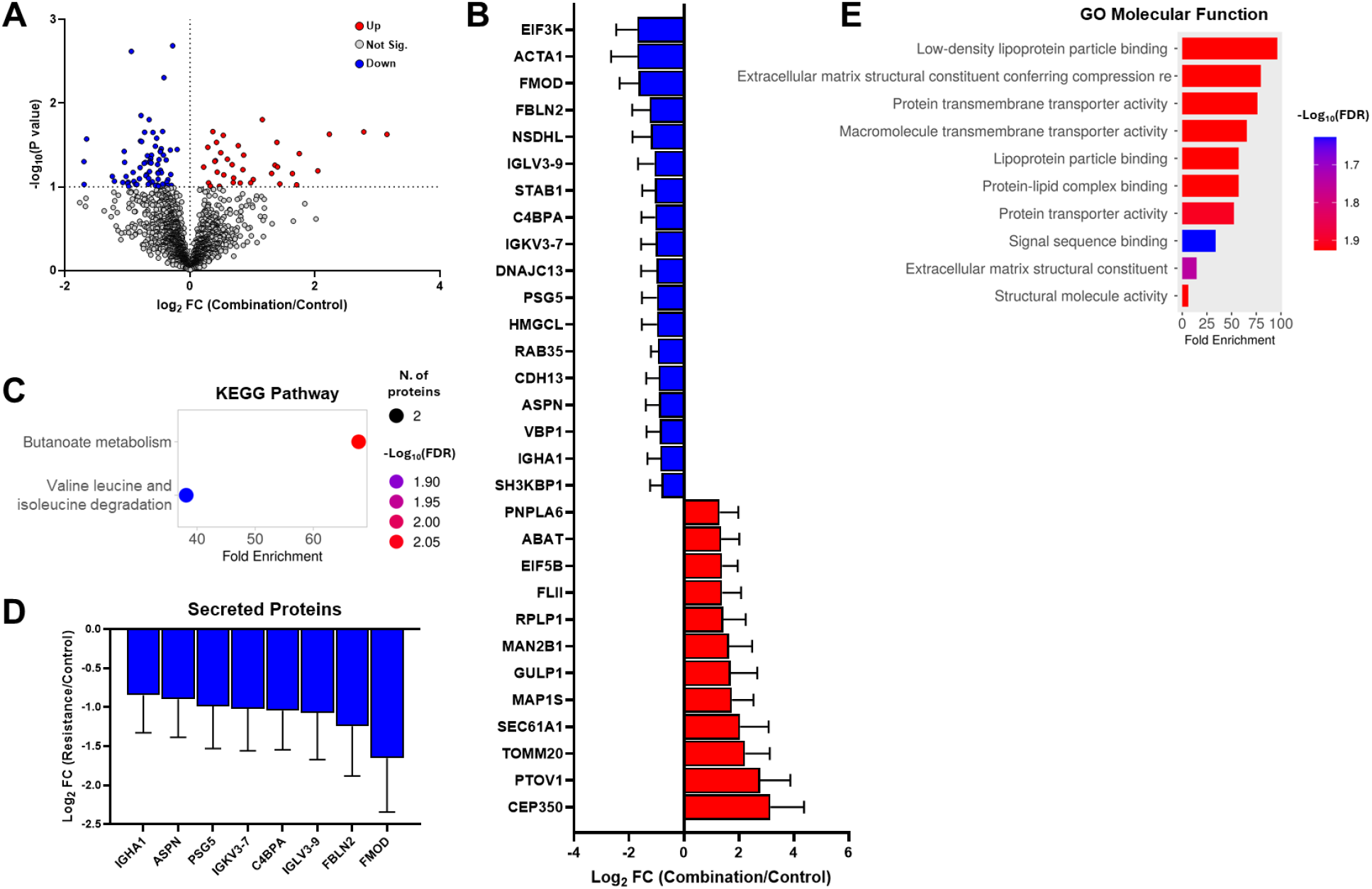
Effects of combination exercise on the placental proteome. Volcano plot of all identified proteins with significantly up- and downregulated proteins colored in red and blue, respectively (A). Differentially abundant proteins in response to combination exercise that are within fold-change parameters (>1.2 and <-0.8) (B). KEGG Pathways derived from the differentially abundant proteins (C). Expression of secreted proteins based on the Protein Atlas (D). GO Molecular Function analysis of differentially abundant proteins in combination group (E).

### Comparison of the effects of different maternal exercise modes on placental proteome

To better understand the impact different maternal exercise modes have on remodeling the placental proteome, we compared differentially abundant proteins identified as having significant and relevant changes in each exercise group relative to the control group (Figure 5A). We observed that only Eukaryotic translation initiation factor 3 subunit K (EIF3K, uniport number: Q9UBQ5) decreased in all three exercise groups compared to control group (Supplementary Figure 1A). Additionally, SH3-domain kinase binding protein 1 (SH3KBP1, uniport number: Q8TBC3) decreased with resistance and aerobic exercise (Supplementary Figure 1B), while lower abundance of Dematin Actin Binding Protein (DMTN, uniport number: Q08495) was common for combination and aerobic exercise groups (Supplementary Figure 1C). Finally, resistance and combination exercise had 7 proteins in common (Supplementary Figure 1D). Collectively, these analyses suggest that maternal exercise alterations to the placental proteome are mode specific. To further examine these mode-specific dependencies, we compared the significant and relevant protein abundance differences arising from exercise group comparisons (Supplementary Figure 2A-C). Interestingly, as shown in Figure 5B, there are no overlapping proteins.

**Figure 5.**
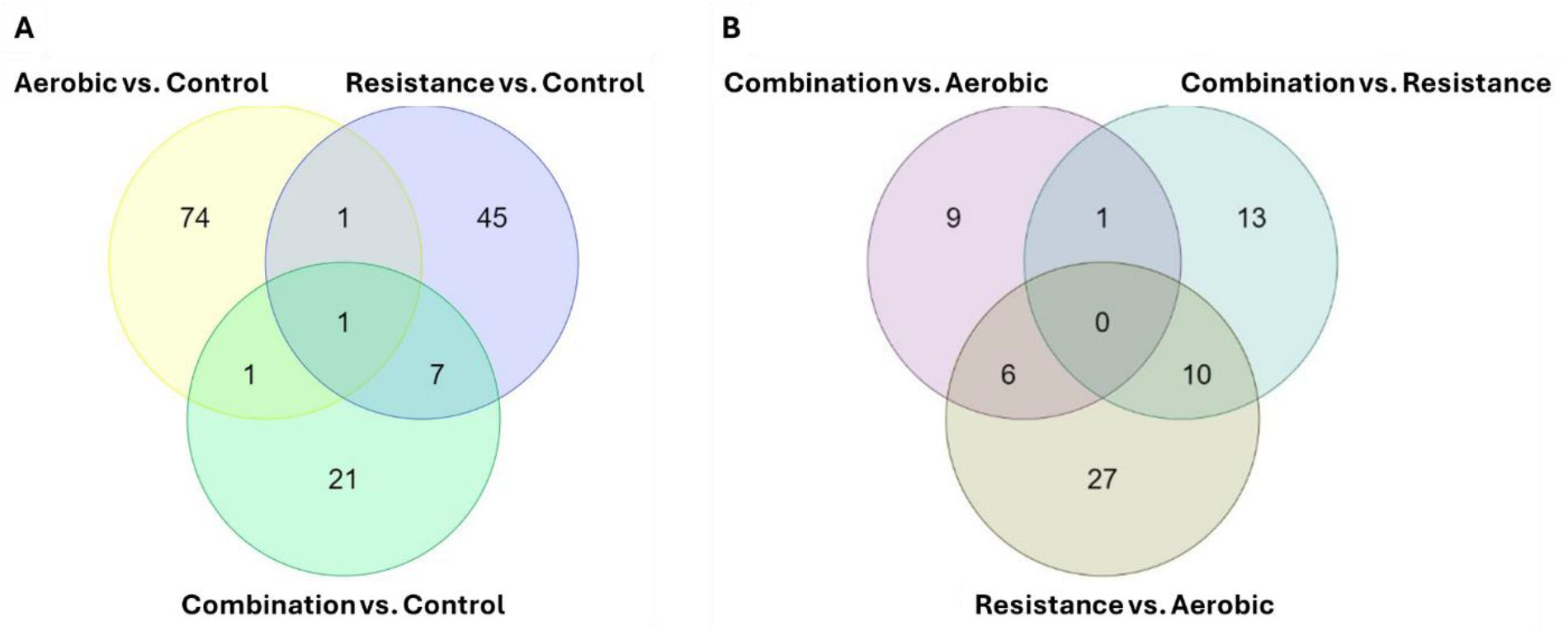
Maternal exercise modes differently affect placental proteome. Venn diagrams of significantly altered proteins in response to different maternal exercise modes (A). Venn diagrams of identified proteins in different modes of exercise (B).

### Correlation of the placental proteome with maternal exercise volume

We performed correlation of 2025 identified placental proteins with metrics of exercise volume. We observed that the average weekly and total pregnancy exercise volume were associated significantly with 374 and 449 placental proteins out of the 2025 identified, respectively (Figure 6A-B). KEGG pathway analysis on the proteins inversely associated with exercise volume corresponded to the TCA cycle, carbon metabolism, and other metabolic pathways. On contrary, proteins positively associated with exercise volumes corresponded with ribosome, spliceosome, protein export and other KEGG pathways. Together these data suggest that maternal exercise volume could play a role in altering placental proteome.

**Figure 6.**
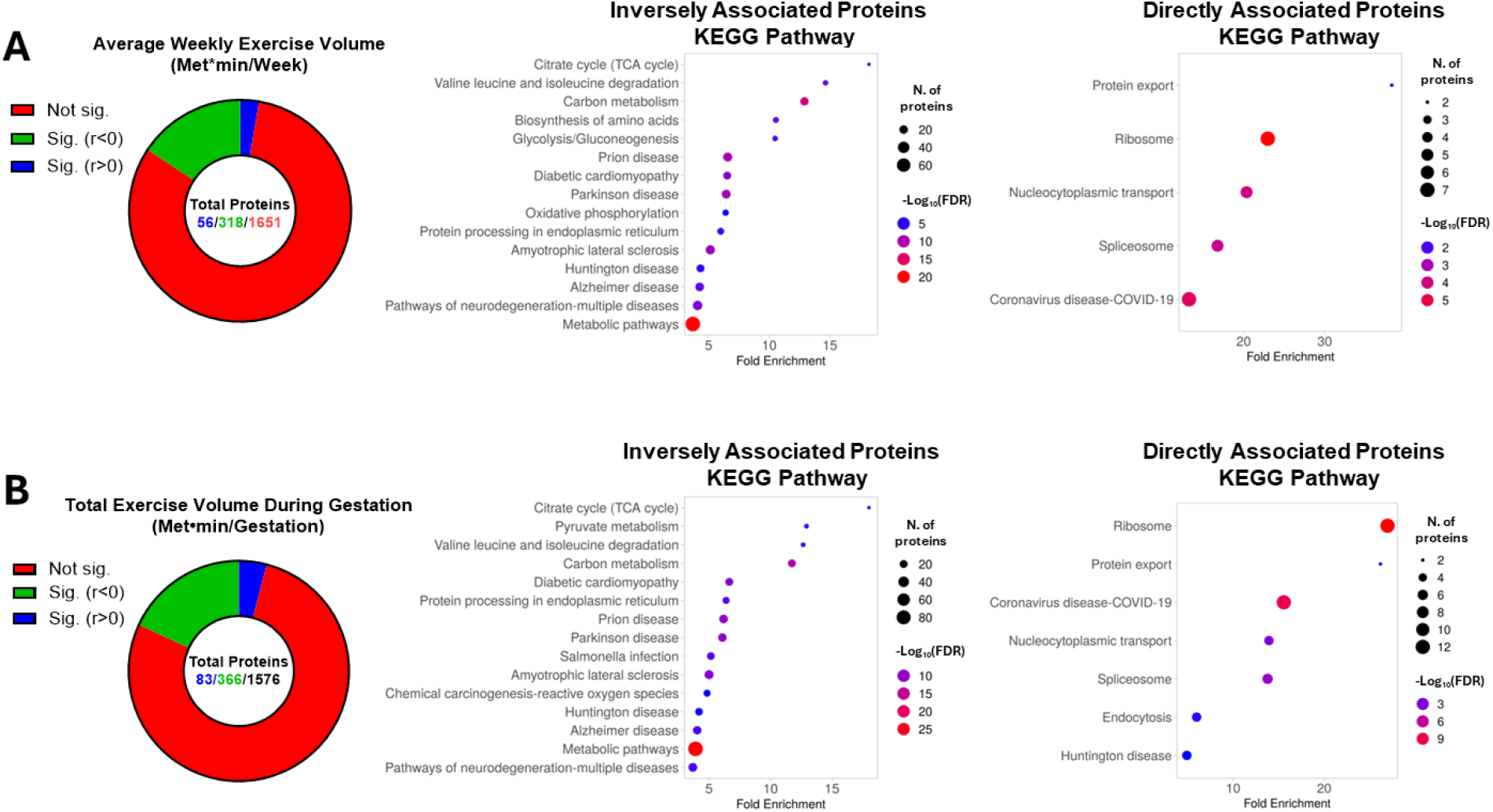
Association of maternal exercise volume with placental proteome. Doughnut charts with the proportion of the placental proteome associated with maternal weekly (A) and gestational (B) exercise volumes (green=significantly inversely associated proteins; blue, significantly directly associated proteins). KEGG pathways to the right of the doughnut charts represent the pathways that significantly correlated proteins correspond to. Pearson correlation, p<0.05. n=40.

## Discussion

The American College of Obstetrics and Gynecology (ACOG) recommends prenatal physical activity as a safe, desirable, and essential element of a healthy lifestyle and pregnancy[38]. They further provide a list of exercises that are deemed safe and beneficial, including walking, dancing, aerobic and resistance exercise, as well as a range of exercise volumes that should be met weekly. Although these recommendations represent an excellent starting point, recent studies suggest that there may be a mode-specific and dose-dependent effect of maternal exercise on maternal and infant health (reviewed in [39]). In this study, we aimed to elucidate the effect of different types of maternal exercise on the placental proteome. This is the first study to demonstrate vastly different adaptations of the placental to aerobic, resistance, and combination exercise, as well as the association of a large portion of the placental proteome with exercise volume. While it is difficult to speculate how the observed remodeling of the placental proteome translates to maternal and infant health, these data certainly lend credence to future studies aiming to explore the distinct effects of different maternal exercise modes and volumes on the mother and infant.

Aerobic exercise is performed at relatively low workload with high repetition, while resistance training consists of low repetition exercise against high workload. Accordingly, these exercise modes differ in their contractile and energetic demands, as well as the physiological adaptations they elicit. Exercise adaptations are in part mediated by the release of bioactive molecules called cytokines. Unique cytokines are released from various organs and tissues, and depending on their tissue of origin are termed adipokines (secreted by adipose tissue), myokines (secreted by muscle), placentokines (secreted by placenta), etc. These molecules orchestrate organ crosstalk and adaptations, and in pregnancy affect placental and fetal development[25,26]. Like the exercise adaptations, the expression of specific cytokines is largely dependent on exercise mode, intensity, and volume. For example, myokine (and adipokine) irisin levels increase with the onset and progression of pregnancy and have been shown to regulate trophoblast differentiation via AMPK activation[40]. Currently, studies have shown conflicting results regarding the impact of different exercise modes on irisin levels in non-gravid populations [41–44]. Such differences in irisin (as well as other cytokines) expression with different exercise modes, could have further implications in gravid exercisers and subsequently placental development. Attesting to this, our data clearly showcases the differences in adaptations of the placenta to different exercise types with minimal overlap in the alterations to the proteome, which could in part be a result of differential cytokine expression profile. Furthermore, it has been observed that maternal exercise modes have differential effects on maternal gestational health[31,45], placental size [6], infant mesenchymal stem cell glucose metabolism [8,12], and infant birth anthropometrics[6].

In addition to the exercise mode, exercise volume has been established as a crucial component of exercise-induced benefits in non-gravid and gravid populations. Here we show that a large part (∼19-23%) of the proteome is associated with metrics of maternal weekly exercise volume, and total volume of exercise performed during gestation. Specifically, some of the pathways associated with exercise volume were related to placental metabolism (TCA cycle, carbon metabolism, metabolic pathways). Interestingly, there was an inverse relationship between exercise volume and TCA cycle enzymes (e.g., PDHB, CS, IDH2, OGDH, SDHA, MDH1), as well as mitochondrial OXPHOS proteins (e.g., Complex I subunits NDUFS1, 4; NDUFA2, 9; ETFA, ETFB, etc.). This is surprising considering that such changes have been associated with maternal obesity and further correspond to lower placental mitochondrial capacity and content [46,47]; however, it is hard to determine whether association with protein abundance translates to changes in placental bioenergetics. Nonetheless, considering the importance of placental energy metabolism in maternal and fetal health and disease, it is imperative to continue investigating the association of exercise frequency, intensity, time, type, and volume (FITT-V) with changes in the placental proteome.

The superoxide dismutase (SOD 1-3) antioxidant enzymes catalyze the scavenging of superoxide radicals to protect various tissues from oxidative stress. Recently placental SOD3 has been shown to mediate the benefits of maternal exercise on mouse offspring glucose tolerance by decreasing oxidative stress-induced methylation of fetal liver genes involved in glucose metabolism [48,49]. More importantly, active compared to inactive women (determined by accelerometers in free-living conditions at mid-gestation) had higher SOD3 mRNA expression in the placenta at parturition. Contrary to these results, when compared to the control group, the aerobic, but not combination or resistance exercise groups, had significantly lower placental SOD3 protein expression (p=0.037, Supplementary Figure 3A). Additionally, we observed an inverse relationship between average weekly and total gestational exercise volume with SOD3 (Supplementary Figure 3B-C), and SOD1, but not SOD2 placental expression (data not shown). These differences are surprising and discrepancies in how exercise volume was assessed (controlled and supervised vs. assessed via accelerometer), what exercise mode was performed, and other factors such as maternal metabolic health could play a role in these differences. In line with that, we observed a significant direct correlation between placental SOD3 and maternal 16-week LDL (p=0.049, r=0.33), non-HDL (p=0.037, r=0.34), and total cholesterol (p=0.051, r=0.32) (data not shown).

Current findings support and extend the growing evidence for promoting exercise during pregnancy. Importantly, these findings highlight the susceptibility of the placental proteome to different maternal exercise modes. Strengths of this study are a prospective, randomized controlled trial study design which provides the strongest evidence for causality. Although, we acknowledge that our sample consisted of “apparently healthy” pregnant women, we had a range of BMI, healthy weight, overweight, and obese, which helps the generalizability of our findings. Further, we did not account for participant dietary habits, or paternal obesity which could be confounding factors in placental adaptations. Future directions should address these limitations and focus on delineating the effects of maternal exercise modes before and during pregnancy on placental, maternal, and infant metabolic health.

In conclusion, this study aimed to assess how the placental proteomic landscape is remodeled in response to supervised modes of ME. Here, we show that placental adaptations to maternal exercise are mode-specific and associated with exercise volume. Finally, these data showcase myriad of altered placental pathways which incite further investigation to determine how these changes affect maternal and infant health.

## Acknowledgments

We thank Lindsey Rossa and Caitlyn Ollmann for assisting with specimen collection and subject recruitment. We thank the subjects for their participation.

