## Supplementary Figure for "Maternal exercise alters placental proteome in an exercise mode-specific manner"

**Disclosures**: The authors have nothing to disclose.

**Funding**: This project was supported by the American Heart Association (18IPA34150006) and NIH (5R01DK129480-01) to L.E.M, and National Institutes of Health grant R01 DK101043 (to L.J.G.), and P30 DK036836 (DRC to Joslin Diabetes Center). Mass spectrometry acquisition and analysis performed in the Brody School of Medicine at East Carolina University's Mass Spectrometry Core was supported in part by the Golden Leaf Foundation and from federal COVID-19 relief funds appropriated to ECU in North Carolina SL 2020-4.

**Clinical Trial:** ClinicalTrials.gov Identifier: NCT03838146 and NCT04805502

**Author Contributions:** F.Jev. conceptualized and wrote the manuscript. F.Jev. and B.W. collected the data. F.Jev., B.W., T.N.Z. managed and analyzed the data. F.Jah., B.W., A.C., D.N.C., J.E.D., S.M., T.N.Z., L.J.G., and L.E.M., revised and confirmed the integrity of the work. All authors have read and agreed to the published version of the manuscript.

**Acknowledgments:** We thank Lindsey Rossa and Caitlyn Ollmann for assisting with specimen collection and subject recruitment. We thank the subjects for their participation.

### Supplementary Figures:

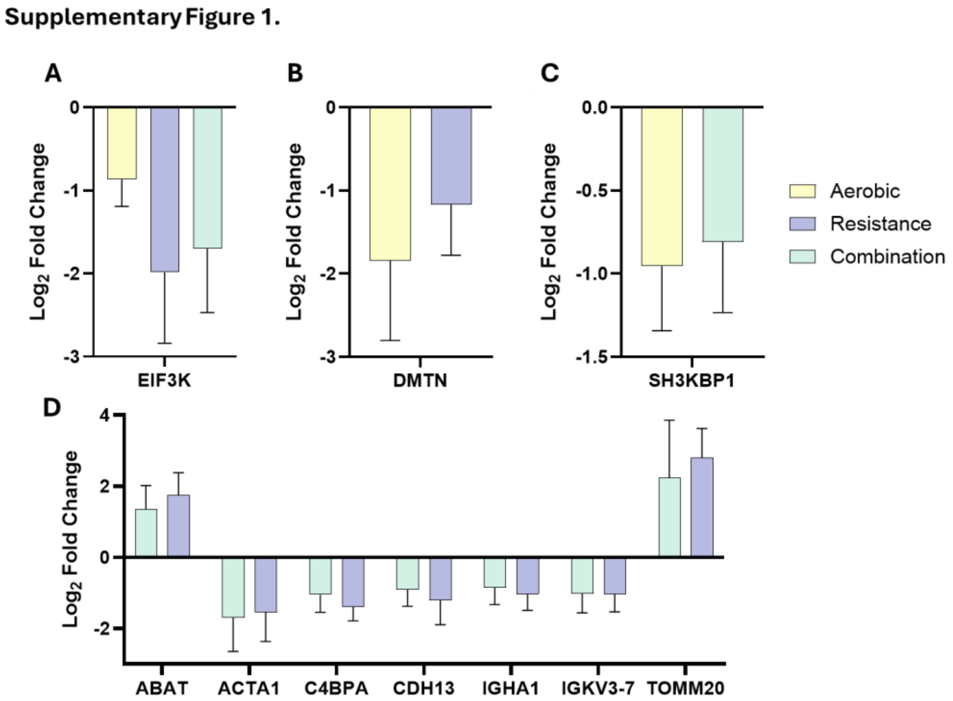


**Supplementary Figure 1.** When compared to the control group, placental proteins were altered by all three groups (A), aerobic and resistance only (B), aerobic and combination (C), and combination and resistance groups (D).

### Supplementary Figure 2.

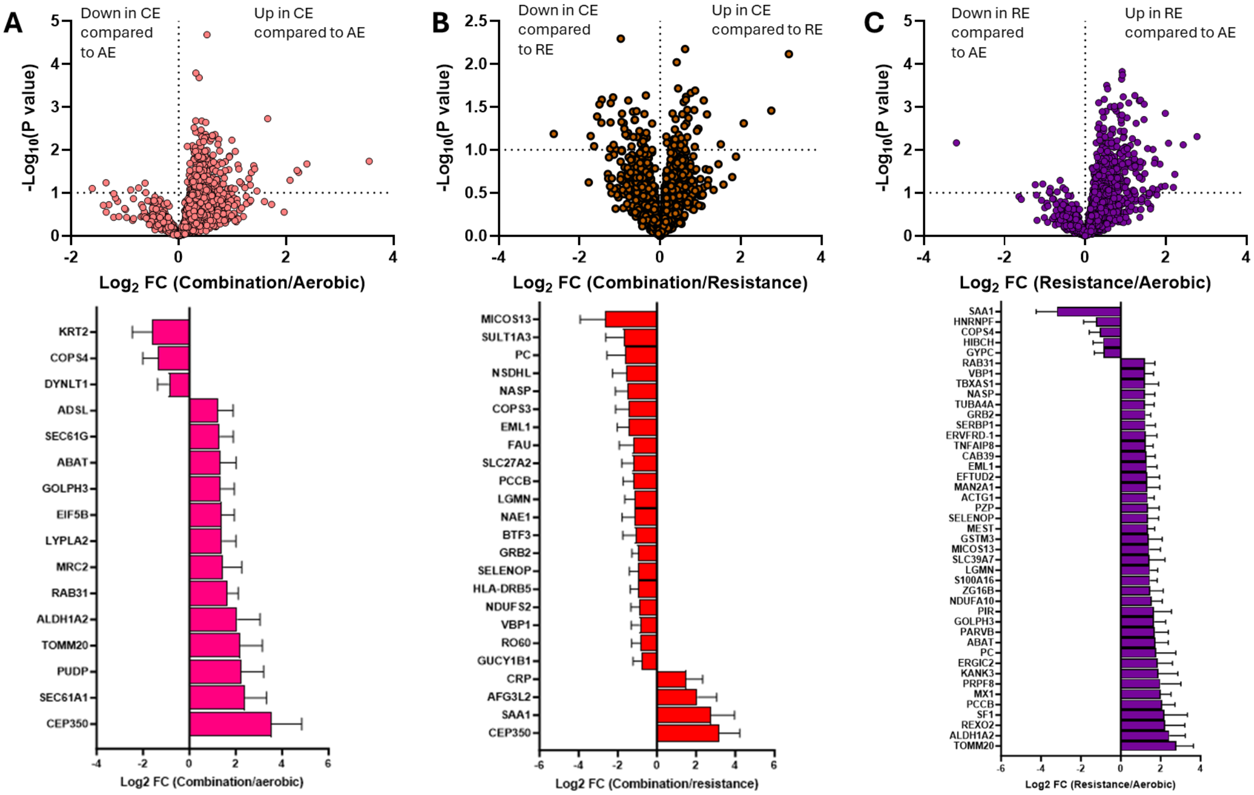


**Supplementary Figure 2.** Vulcano plots with differentially expressed proteins between exercise groups with significant proteins within the fold-change parameters shown below.

### Supplementary Figure 3.

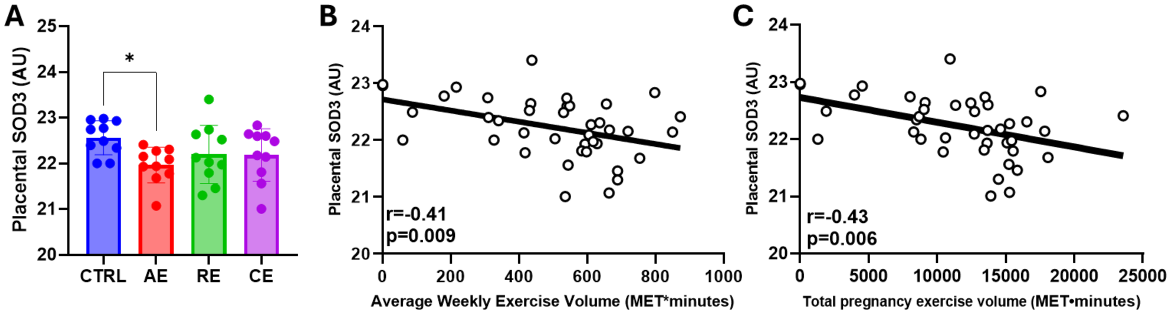


**Supplementary Figure 3.** Placental sodium dismutase 3 (SOD3) expression across groups (A). Pearson's correlation between SOD3 and average weekly exercise volume (A) and total exercise volume during pregnancy (C).
